# Enhanced Pipeline ‘MetaGaAP-Py’ for the Analysis of Quasispecies and Non-Model Microbial Populations using Ultra-Deep ‘Meta-barcode’ Sequencing

**DOI:** 10.1101/171520

**Authors:** Christopher Noune, Caroline Hauxwell

## Abstract

A pipeline developed to establish sequence identity and estimate abundance of non-model organisms (such as viral quasispecies) using customized ultra-deep sequence ‘meta-barcodes’ has been modified to improve performance by re-development in the Python programming language. Redundant packages were removed and new features added. RAM and storage usage have been optimized to facilitate the computational speeds though coding optimizations and improved cross-platform compatibility. However, computational limits restrict the approach to barcodes spanning a maximum of 30 polymorphisms. The modified pipeline, MetaGaAP-Py, is available for download here: https://github.com/CNoune/IMG_pipelines

## 1. Introduction

The ‘Meta-Barcoding Genotyping and Abundance Pipeline’ (MetaGaAP) was developed to identify and estimate abundance of strain variants within non-model populations by identification and ultra-deep sequencing of custom ‘meta-barcodes’ and comparison with a database of all possible polymorphisms generated from the sequence data, which was then validated through analysis of quasispecies within baculovirus isolates [1-3]. This approach facilitates analysis of viral quasispecies for which standard ‘barcodes’ sequence databases are not readily available. Since the original release on GitHub, the limits on cross-platform compatibility, the large number of dependencies, the high computation capacity required and reliance on Bash and R programming languages were found to limit performance. These were addressed by redevelopment in Python.

## 2. Method

The Python version 3.6 programming language was selected as a versatile, general purpose, high-level non-compiled language (interpreted language) which is backwards compatible with all versions of Python 3. The pipeline was fully re-coded using the Anaconda 3.6.1 and the Spyder 3.1.4 integrated developer environments [4,5] to ensure no redundant Bash or R code would be carried over.

A core set of dependencies were retained: The Burrows-Wheelers Aligner (version 0.7.15 or above), Samtools (version 1.3 or above), the Genome Analysis Toolkit (version 3.6 or above), fastx-toolkit (0.0.14 or above), Picard-tools (version 2.9 or above), Oracle Java 1.8, mawk (version 1.3.3), Sed (version 4.2 or above) and Biostars175929 that is now included as a pre-compiled version [6-12]. The number of dependencies was reduced: BBmap renamer and duplicate sequence removal tools [13], kentUtils [14], Zenity, and the R coded back-end scripts Subset_Stats.R and Seq_List.R, were discarded and replaced by pure Python implementations coded directly in the source code (Table 1). The implemented Python packages (with the exception of Biopython) are natively-installed with Python 3.6 without the need to write a separate installation script.

**Table 1:** Python implemented dependencies in the revised pipeline

| Package/Library | Function |
| --- | --- |
| TKinter | Implements a simple graphical user interface for file selection. |
| Biopython [15] | A Python library to manipulate fasta sequences. |
| The Python Data Analysis Library (Pandas) | Removes the need to use R code. |
| Multiprocessing | A standard Python library to implement multi-threading. |
| Sys | Captures operating system type i.e. Windows, Linux or Mac. This package was implemented to fix an issue in multi-threading on a Linux system. However, in cases where multi-threading doesn't work, the duplicate removal step will default to a single thread. |
| Garbage Collection | Optimises RAM utilisation. |
| Getpass | Captures user information. |
| OS | Basic Python functions which allow Python to communicate with the operating system. |
| Subprocess | Python functions allowing for the execution of non-Python packages. |

New features were implemented to allow for different analysis types and behind the scenes coding enhancements to improve computational efficiencies (Table 2). To distinguish the original MetaGaAP from the new Python implementation, the original pipeline was renamed as MetaGaAP-Legacy and the new Python implementation named MetaGaAP-Python (MetaGaAP-Py)

**Table 2:** New features added and coding enhancements

| Feature/Enhancement | Function |
| --- | --- |
| Multi-reference, multi-sample analysis | Multiple samples with different reference sequences can be analysed at the same time, e.g. two different 'barcodes' or viruses. |
| Single-reference, multi-sample analysis | Multiple samples using the same reference sequence can be analysed, e.g. time-course analysis |
| Automatic directory creation | Directories are created at the same time as processing to reduce user interactions needed to select output directories. |
| RAM optimised, multi-threaded processing, Python-native duplicate sequence removal [16] | Multi-threaded duplicate sequence removal has been implemented to reduce computational time. In some-cases it may default to a single thread. Optimises RAM usage by creating a new database at the same time as duplicate removal facilitates use on systems with lower than the recommended 8 gigabytes. |
| Sequence combinations database compression by converting multi-line fasta sequences to a single-line fasta sequence [17] | Internal testing has found that the database memory footprint reduces when converted from a multi-line fasta to a single-line fasta file i.e. a single fasta sequence per line rather than a wrapped fasta sequence. |
| Automatic average sequence length and maximum read depth calculation | Automatically tells the HaplotypeCaller the maximum read depth for identification of polymorphisms and the Biostars175929 tool the sequence length required to produce the combinations database. |

## 3. Results and Conclusions

Porting to Python and improved package selection has resulted in a highly-refined pipeline with an optimized workflow. Furthermore, the reduction in required dependencies and coding in a cross-platform compatible language enables execution in Mac OS X, the Microsoft Windows 10 Linux Subsystem and all Linux distributions.

The introduction of multi-reference, multi-sample analysis expands the application to analyse sequence sets from multiple samples with different reference sequences at the same time, e.g. multiple samples using two different ‘barcodes’. Single-reference, multi-sample analysis enables analysis of sequences from multiple samples using the same reference sequence such as time-course analysis of viral quasispecies.

Some weaknesses persist. The implemented duplicate sequence removal is dependent on the number of cores in the user’s central processing unit, on whether the storage unit is a solid-state drive or a mechanical hard-disk drive, and on the size of dataset: the pipeline is functionally limited to analysis of a maximum of 30 polymorphisms across the barcode region due to the lack-of multi-threading within the Biostars175929 tool and memory requirements to store large databases. In addition, fastq files and sample names need to be re-specified when completing the final mapping stage and calculating abundance result as part of a single-reference, multi-sample analysis.

Overall the optimisations, newly implemented features, and reduced dependency requirements facilitate the use of MetaGaAP-Py, resulting in a less computationally demanding and more streamlined user interface that can be applied to generation and application of customised sequence ‘barcodes’ and libraries for identification and quantification of quasispecies variants in non-model populations, for which standard sequence ‘barcodes’ and public sequence databases are not available.

## Acknowledgments

This work was funded by the Queensland University of Technology (QUT), the Cotton Research Development Corporation and an Australian Government Research Training Program Scholarship. We would like to acknowledge the support of the Invertebrate & Microbiology Group at QUT for their assistance. Some of the data reported in this paper was obtained at the Central Analytical Research Facility (CARF) operated by the Institute for Future Environments (QUT). Access to CARF is supported by generous funding from the Science and Engineering Faculty (QUT).

## Author Contributions

Christopher Noune conducted the programming. Both authors contributed equally to the development of concepts and applications, and to the writing of the manuscript.

## Software Availability

MetaGaAP-Py is available for download at https://github.com/CNoune/IMG_pipelines.

